# Significant competitive dominance in mid-latitude European plant communities

**DOI:** 10.1101/056309

**Authors:** José A Capitán, Sara Cuenda, Alejandro Ordóñez, David Alonso

## Abstract

Understanding the main determinants of species coexistence across space and time is a central question in ecology. However, ecologists still know little about the scales and conditions at which biotic interactions matter and their interplay with the environment to structure species assemblages. Here we develop ecological theory to analyze plant distribution and trait data across Europe and find that plant height clustering is related to evapotranspiration and gross primary productivity. Our analysis suggests competitive dominance as a plausible mechanism underlying community assembly patterns over these continental scales. In particular, we find a clear signal of plant-to-plant competition in mid-latitude ecoregions, where conditions for growth (reflected in actual evapotranspiration rates and gross primary productivities) are optimal. Under severe conditions, either climate is too harsh and overrides the effect of competition or other interactions play a relevant role. Our approach bridges the gap between modern coexistence theory and large-scale species distribution data analysis.

---

Classical coexistence theory [1, 2] assumes that the more similar two species are in their niche requirements, the more strongly they will compete over shared resources, an idea that can be traced back to Darwin [3]. Ever since, competition-similarity hypotheses have been at the front line of theoretical explanations for species coexistence [2, 4], exploring community assembly based on phylogenetic or functional similarity. This framework predicts that large species differences should be selected during community assembly to reduce competition. Therefore, trait and/or phylogenetic overdispersion have often been regarded as signatures of competitive interactions. However, progress in our understanding of how species differences influence the outcome of competitive interactions [5, 6] shows that this theoretical framework is too simplistic because it disregards the balance between stabilizing and equalizing species differences [5]. Stabilizing mechanisms are based on trait differences that cause species to be limited more by their own con-specifics than by their competitors, favoring species when they drop to low densities, which, in turn, promotes species coexistence. Equalizing mechanisms, by contrast, promote species dominance over potential competitors. In the absence of stabilizing species differences, superior competitors would drive other species to extinction through competitive exclusion. In communities controlled by equalizing mechanisms, species with similar trait values should be selected through competitive dominance, resulting in high levels of trait clustering even in the absence of environmental filtering [6]. This theoretical framework suggests that significant trait clustering at local sites may be a fingerprint of biotic (competitive) interactions controlling the composition of ecological communities.

Accurately separating the effect of biotic interactions from environmental filters as structuring agents of community assembly is not trivial. Despite the undeniable success of species distribution models [7], there is an increasing recognition of the need for simple, process-based models to make robust predictions that help understand species responses to environmental change [8]. However, we still lack clear evidence for the role of biotic interactions in shaping species assemblages at large spatial scales. Studies based on species randomization models have attempted to separate the outcomes of competitive exclusion and environmental filtering by assuming the competition-similarity hypothesis as a given [9, 10]. Empirical studies, while they may be able to independently assess environmental stress and species competitive abilities, are often limited to small community sizes [11] or restricted to single habitats [12]. Very few studies have explored the idea of competition as a driver of community assembly across biogeographic regions [13, 14]. Here we report the results of the first macro-ecological study, based on theoretical predictions from modern coexistence theory [5, 6], aimed at separating the combined effects of biotic (competition) and abiotic (environmental) factors shaping plant community assemblages at large geographical scales.

Light and water availability impose significant limitations on gross primary productivity which is reflected in actual evapotranspiration rates [15]. These two resources vary at regional scales, placing strong, but opposing constraints on how tall a plant can grow within the limits of structural stability [16, 17, 18, 19]. In addition, plants have to compete strongly for these resources. The resulting plant height is a trait that reflects the ability of the individual to optimize its own growth within these regional environmental and biotic constraints (see [20, 21] and references therein). We analyzed presence-absence matrices of floral taxa across different European ecoregions to determine if competitive ability (reflected in maximum stem height) could help explain assemblage patterns at local scales across gradients of relevant environmentally-driven factors such as evapotranspiration. We examined how macro-ecological plant data match up to theoretical predictions generated by a synthetic, stochastic framework of community assembly [22, 23]. Competition between hetero-specifics was measured by signed height differences, so that clustering could emerge by competitive exclusion of sub-dominant species. We find large fractions of local communities where clustering in maximum stem height is significant at intermediate latitudes, coinciding with a mid-latitude peak in evapotranspiration rates. Across Europe, actual evapotranspiration is lower at more southern latitudes (due to reduced precipitation levels) as well as at more northern latitudes (due to colder temperatures and low levels of sunlight). Species trait clustering is significant only in a latitudinal band where environmental constraints to plant growth are weaker, suggesting that it is only in these mid-latitude ecoregions that a clear signature of competitive dominance can be found in species assemblage patterns.

## Results

### European plant ecoregions

Plant community data were drawn from Atlas Florae Europaeae [24]. The distribution of flora is geographically described using equally-sized grid cells (~ 50 × 50 km) based on the Universal Transverse Mercator projection and the Military Grid Reference System, see Fig. 1. Each cell was assigned to a dominant habitat type based on the WWF Biomes of the World classification [25], which defines different ecoregions, i.e., geographically distinct assemblages of species subject to similar environmental conditions. We consider each cell in an ecoregion to represent a species aggregation (which we name ‘local community’. We also refer to ecoregions as ‘metacommunities’. A total of 3233 plant taxa were extracted from data sources.

Each species in an ecoregion was characterized by its maximum stem height *H*, an eco-morphological trait that relates to several critical functional strategies among plants. It represents an optimal trade-off between the gains of accessing light [16, 17], water transport from soil [18, 19], and sensitivity to biomass loss from mechanical disturbances [26], and is therefore a good proxy for competitive interactions at the local level under light- or water-limited conditions [21, 27]. Thus, within the limits of plant structural stability, stem height results from the interplay of two opposing forces: competition for light and competition for water resources. In situations of low water availability, the cost of transport water to height increases [18, 19] and hydric stress tends to make individuals shorter. On the contrary, if water resources are abundant, individual plants compete for light and taller individuals are more favored than neighbors [16, 17]. In front of these opposing constraints, depending on environmental conditions, plants optimize their strategy, which, therefore, must be reflected in the selection for large (competition for light) or small (competition for water) individuals at local scales [21], see Fig. 2a. Height values were compiled from the LEDA database [28] (Methods).

**Figure 1.**
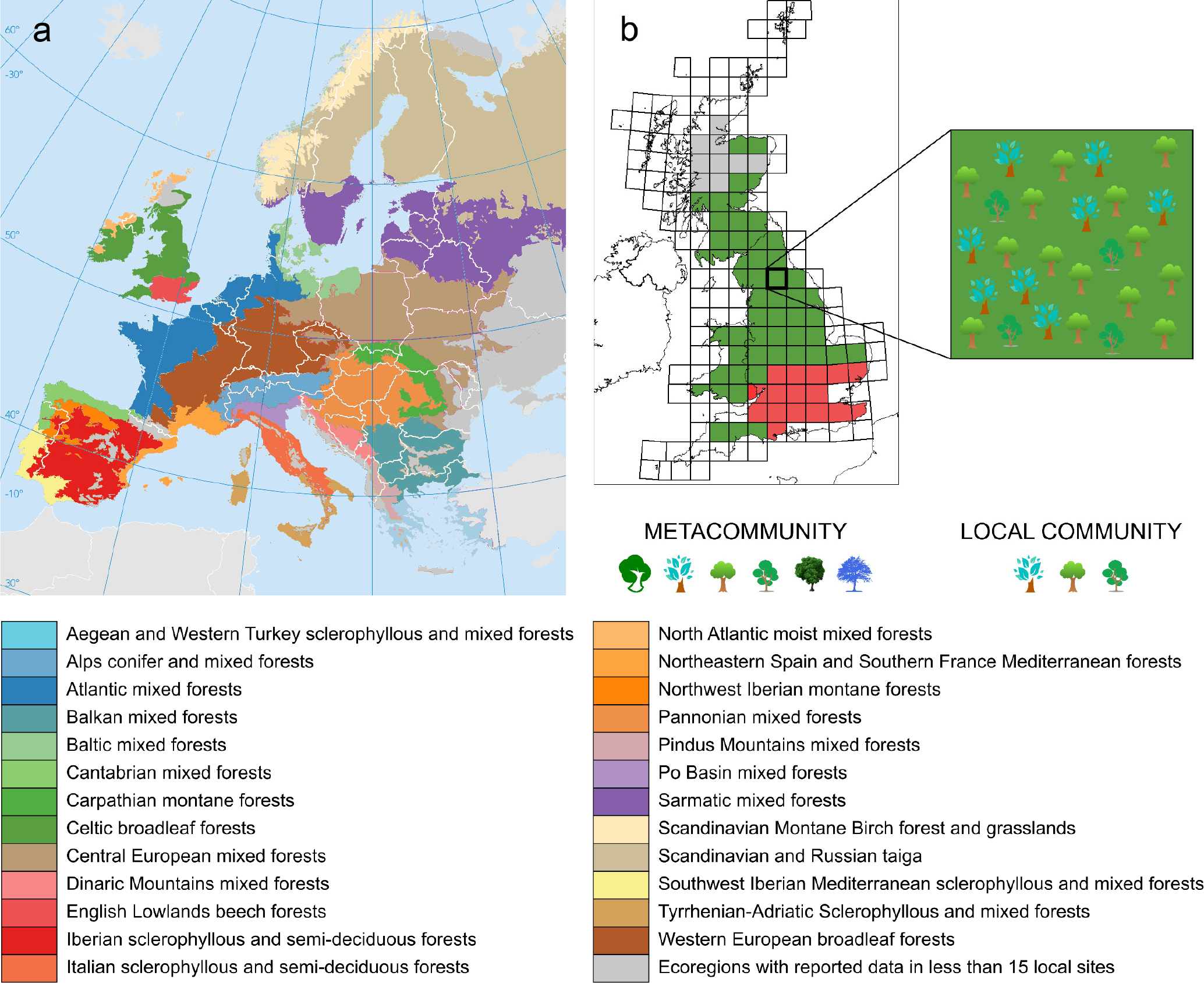
Geographical description of plant data across European ecoregions. **a**, 25 different habitats covering most of Europe are shown in the map and listed below. Ecoregions are regarded as metacommunities comprising all plant species observed in that region. **b**, The Military Grid Reference System divides ecoregions in grid cells, each one considered as a local community formed by a species sample of the metacommunity.

Competition for light is asymmetric [17, 29]. Given that the costs of transporting water in situations of hydric stress increase with height [30], competition for water can also be regarded as asymmetric. Therefore, species competitive dominance was measured by signed height differences, 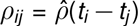, where *t_i_* are height values standardized across metacommunities and are sorted in increasing order (Methods). This choice assumes the subsequent selection for low-trait valued species (Fig. 2b). Alternatively, the opposite choice 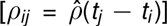 implies that large traits will be over-represented in local communities (see Fig. 2b and Supporting Information, Secs. S2.3 and S3.2). The scale factor 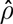 measures the ratio between inter- *vs*. intraspecific competition. For all the species reported in an ecoregion, we formed a competition matrix with the pairwise *ρij* values. The advantage of having these values represent trait differences between pairs of species is that any trend in competition strengths can be immediately translated into patterns of functional clustering or overdispersion. For a metacommunity with *S* species, we calculated the average competitive strength as 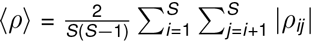.

**Figure 2.**
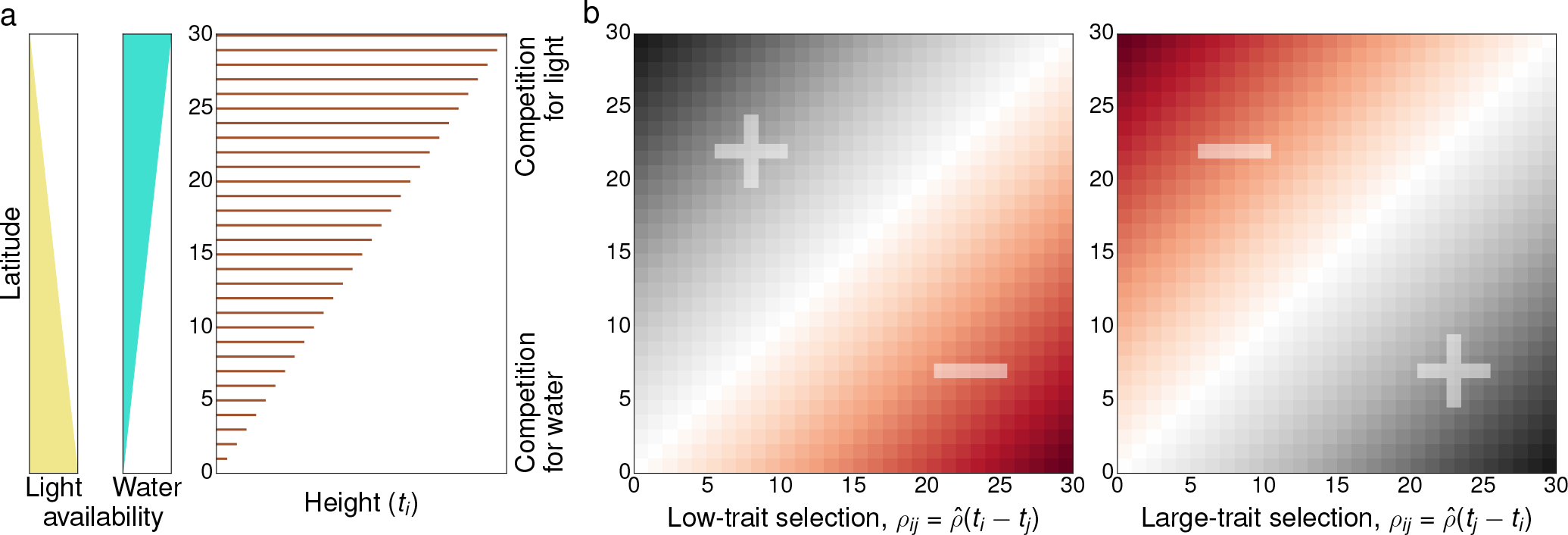
Conceptual framework for height as a trade-off between accessing light and water resources. **a**, Here we illustrate the influence of light and water availability profiles in the resulting plant height. To simplify we assume that light access and water availability exhibit opposing latitudinal gradients. Hence, plants in southern latitudes will be selected by competition-for-water mechanisms, which tend to give advantage to smaller individuals, whereas competition for light (favoring tall plants) will dominate over northern latitudes. This conceptual framework is consistent with the height profile represented in the third panel (the vertical axis stands for the species index *i*). **b**, According to the trait profile, competitive dominance matrices can be chosen either as 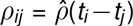 or as 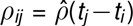. Given that a competitive strength *ρij* < 0 increases the growth rate of species *i* (Supporting Information, Sec. S2.1), the two choices lead to the selection of low- and large- trait values, respectively. Positive (negative) matrix entries have been marked in black (red). Rows associated to dark red areas indicate the species indices that will be preferentially present in local communities.

### Two predictions from theory

Community assembly results from the interplay of four fundamental processes: speciation, ecological drift, dispersal, and selection acting across space and time [31]. New species are added to local communities through speciation and/or immigration, species abundances are then shaped by stochastic births and deaths (ecological drift), ongoing dispersal and differential growth driven by species interactions and/or specific adaptations to local environment (selection). Our stochastic approach disregards speciation, does not account for environmental factors and assembles communities based only on dispersal, ecological drift and asymmetric competition processes. The community model is defined by four parameters representing elementary processes, namely: *α*^+^ (local birth), *α*^−^ (local death),*µ* (immigration of newcomers to local communities), and *K* (interspecific competition is proportional to *αρij*/*K*, where *K* is interpreted as a carrying capacity and α = |α^+^ – α^−^|). In model simulations, trait values *t_i_* are drawn from a Gaussian distribution with zero mean and variance *σ*^2^,and trait differences are transformed to competitive strengths afterwards (Methods).

The stochastic dynamics predicts the identities of species selected by competition in local communities, as well as the observed local diversity relative to the metacommunity richness (we refer to this ratio as ‘coexistence probability’ and denote it as p_c_, see Methods). We used the variability in coexistence probability as a function of the average competitive strength 〈*ρ*〉 and the distribution of pairwise competitive strengths observed in real communities to test how closely they confirm the predictions of the stochastic community assembly model.

**Figure 3.**
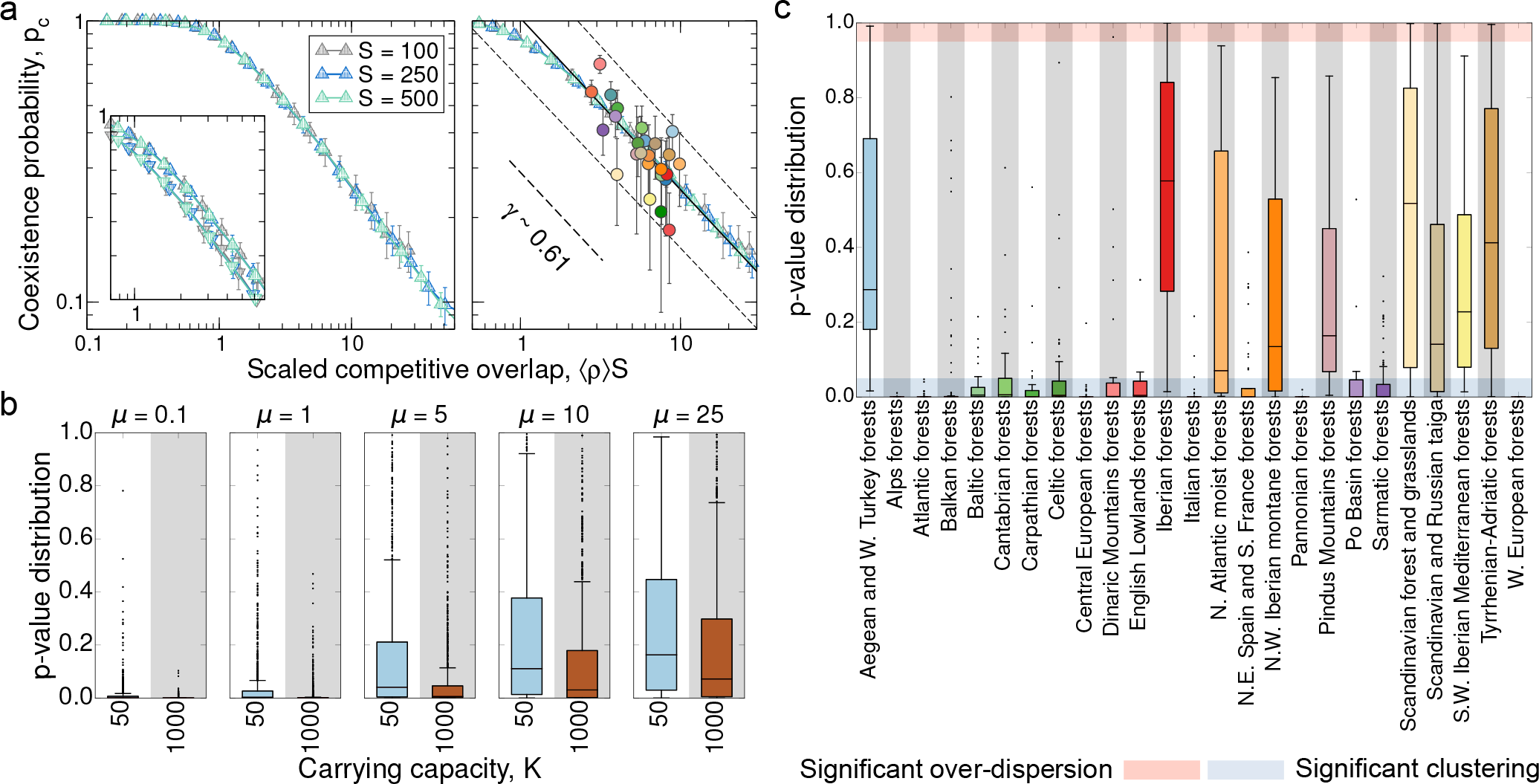
Two theoretical predictions tested against data. **a**, Left: Average fraction p_c_ of coexisting species in local communities as a function of the scaled competitive strength, 〈*ρ*〉*S*. Simulation parameters are *α*^+^ = 50, *α*^−^ = 0.1, *µ* = 5, *σ* = 0.2 and *K* = 50 (down triangles, inset) or *K* = 1000 (up triangles). Right: The model predicts a power-law decay *p*_c_ ~ (〈*ρ*〉*S*)^−γ^ regardless of the metacommunity size *S*, which permits fitting a power law to data (*r*^2^ = 0.52, *p* < 10^−3^, 95% prediction intervals shown). In order for model curves to match empirical data, we need to choose 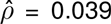 in the calculation of *ρij* (Supporting Information). **b**, Model randomization tests for different immigration rates and carrying capacities; here 〈*ρ*〉 = 0.06 and *S* = 100 (Supporting Information, Sec. S2.5). The closer the distribution is to 0, the larger is the fraction of cells where trait clustering is significant. For parameter values that fit data in the low immigration regime (µ ~ 5, *µ*/*αK* ≪ 1, panel **a**), the model indicates a clear signature of clustering. c, Empirical randomization tests. The majority of ecoregions are consistent with model predictions as the distributions (Tukey boxplots) lie in the 5% range of significant clustering. Color codes for data in panels **a** and **c** match codes in Fig. 1.

A first theoretical prediction from the model is that, as competition increases, coexistence probability shows a power-law decay whose exponent is controlled by the immigration rate *µ* (Supporting Information, Fig. S4). In particular, the curves for different metacommunity sizes collapse when represented as a function of the competitive strength scaled by the metacommunity richness *S*, *p*_c_ ~ (〈*ρ*〉*S*)^−γ^ (see Fig. 3a and Supporting Information, Sec. S2.3, for a theoretical derivation of the curve collapse). This collapse eliminates the variability in S, so that empirical coexistence probabilities, which arise from different metacommunity sizes, can be fitted together (Fig. 3a).

To test the significance of competitive dominance in local communities, we generated a second model prediction applying randomization tests to model communities. Null models for community assembly [9] compare the properties of actual communities against random samples of the same size extracted from the metacommunity. This approach assumes that local communities are built up through the independent arrival of equivalent species from the metacommunity [32, 33] regardless of species preferences for particular environments or species interactions (see Supporting Information, Sec. S2.5, for details). Local species richness is determined by inherent site-level differences in the ability to harbor different numbers of species [34]. Our randomization tests were based on average competitive strengths observed in local communities, which are compared to random samples of the same size drawn from the metacommunity. Observed trait differences can be, on average, significantly larger than the average measured for random metacommunity samples, signifying that average trait differences in local communities are larger than expected at random. On the other hand, empirical trait differences can be significantly smaller than those obtained for random metacommunity samples. In either case the null hypothesis (i.e., local communities are built as random assemblages from the metacommunity) can be rejected, which implies clear signals of ‘significant trait overdispersion’ or ‘significant trait clustering’, respectively. Applying the statistical test to every local community in a metacommunity leads to a distribution of *p*-values (Methods).

Fig. 3b shows the distribution of *p*-values observed when simulated local communities are compared to metacommunity samples (Supporting Information, Sec. S2.5). The model predicts high levels of trait clustering for low immigration rates and high carrying capacity values (Fig. 3b), as well as the selection for low/large-trait valued species (depending on how *ρij* is defined, see Fig. 2 and Supporting Information, Sec. S2.3). When immigration increases, local communities show a broad distribution of *p*-values. At the highest immigration rates, model communities are essentially random samples of the metacommunity since immigration overrides competition in this regime [35]. Fig. 3a provides a hint of the range in which model parameters best fit real data. For a realistic fit, the exponent of the empirical power law is obtained for *µ*/*α* ~ 0.1 individuals per generation. Since plant communities operate in a low-immigration regime, the non-dimensional immigration rate λ = *µ*/*αK* must satisfy λ = 0.1/*K* ≪ 1, hence the carrying capacity must be large. In a regime of low immigration rate and high carrying capacity, which best fits empirical coexistence probabilities, the model predicts a significant degree of species clustering (Fig. 3b).

### Testing model predictions against data

Regarding the first prediction, we found a significant correlation between coexistence probability and the scaled competitive overlap based on empirical data (*r*^2^ = 0.52, *p* < 10^−3^, Fig. 3a). Apparently, a model driven solely by dominant competitive interactions reliably predicts the average richness of local plant communities across different ecoregions. In addition, this theoretical prediction allowed an indirect estimation of the relative importance 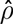 of inter- *vs*. intraspecific average effects: the average ratio of inter- to intraspecific competition strength is about 4% (see Supporting Information, Sec. S2.1, for details on the estimation procedure).

On the other hand, we calculated *p*-values for randomization tests applied to all local communities in each ecoregion, which represent the empirical metacommunity distribution of *p*-values (Fig. 3c). At the parameter values that make plant data consistent with the first prediction, our model predicts significant trait clustering (see Fig. 3b). When this prediction is compared with empirical data, we observe that some ecoregions fit best with our model, based on competitive dominance, while others clearly do not. In addition, no ecoregion is consistent with trait overdispersion (Fig. 3c). We have also conducted randomization tests based on height values, not differences, to check whether the clustering observed in local communities is due to the selection for large or low trait values (Fig. 2). In ecoregions where height clustering is significant, we obtain consistent signatures of small plant selection (see Fig. S8 in Supporting Information and Sec. S3.2). Therefore, we conclude that local community assembly in those ecoregions is plausibly driven by competitive differences biased towards selection for smaller plants.

### Ecoregion clustering and actual evapotranspiration rates

In order to better quantify the propensity of an ecoregion to exhibit clustering in maximum stem height, we define a clustering index *q* for an ecoregion as the fraction of its local communities that lie within the 5% range of significant clustering (randomization tests yield *p*-values smaller than 0.05 for those cells). An ecoregion for which significant clustering is found in most of its local communities will tend to score high in the *q* index. We examined how the clustering index varied across the continent (geographical location of ecoregion centroids) as well as with actual evapotranspiration (Fig. 4). Evapotranspiration maps were obtained from data estimated through remote sensing [36]. Water availability acts as a factor limiting plant growth at geographical scales, and correlates with gross primary productivity [15], see Fig. 4d. Therefore, for a given region, mean annual evapotranspiration is a distinct measure of environmental constraints on plant growth [15]. Panels a and b of Fig. 4 show a clear latitudinal trend: there is an intermediate range of ecoregion latitudes where both clustering indices and evapotranspiration are large, indicating that evapotranspiration measures can robustly predict clustering indices (Fig. 4c). More importantly, since evapotranspiration is a powerful proxy of environmental constraints on plant growth, this clustering in maximum stem height appears to be strongest at ecoregions less limited by environmental conditions. As environments become harsher and less optimal for plant growth, these clustering patterns disappear. This is particularly true for the severe climatic conditions characteristic in the Mediterranean (with erratic rainfall, limited water availability and drought), as well as of boreal zones (with low radiation incidence and cold temperatures). The harshness of these conditions likely override the effects of competition, and other processes such as species tolerances and facilitation [37] may be critical community drivers at climatic extremes. According to model predictions, the overall clustering patterns found at middle-range latitudes are consistent with species competitive dominance driven by height differences, resulting in a local over-representation of competitive dominant species characterized by relatively lower maximum stem heights.

**Figure 4.**
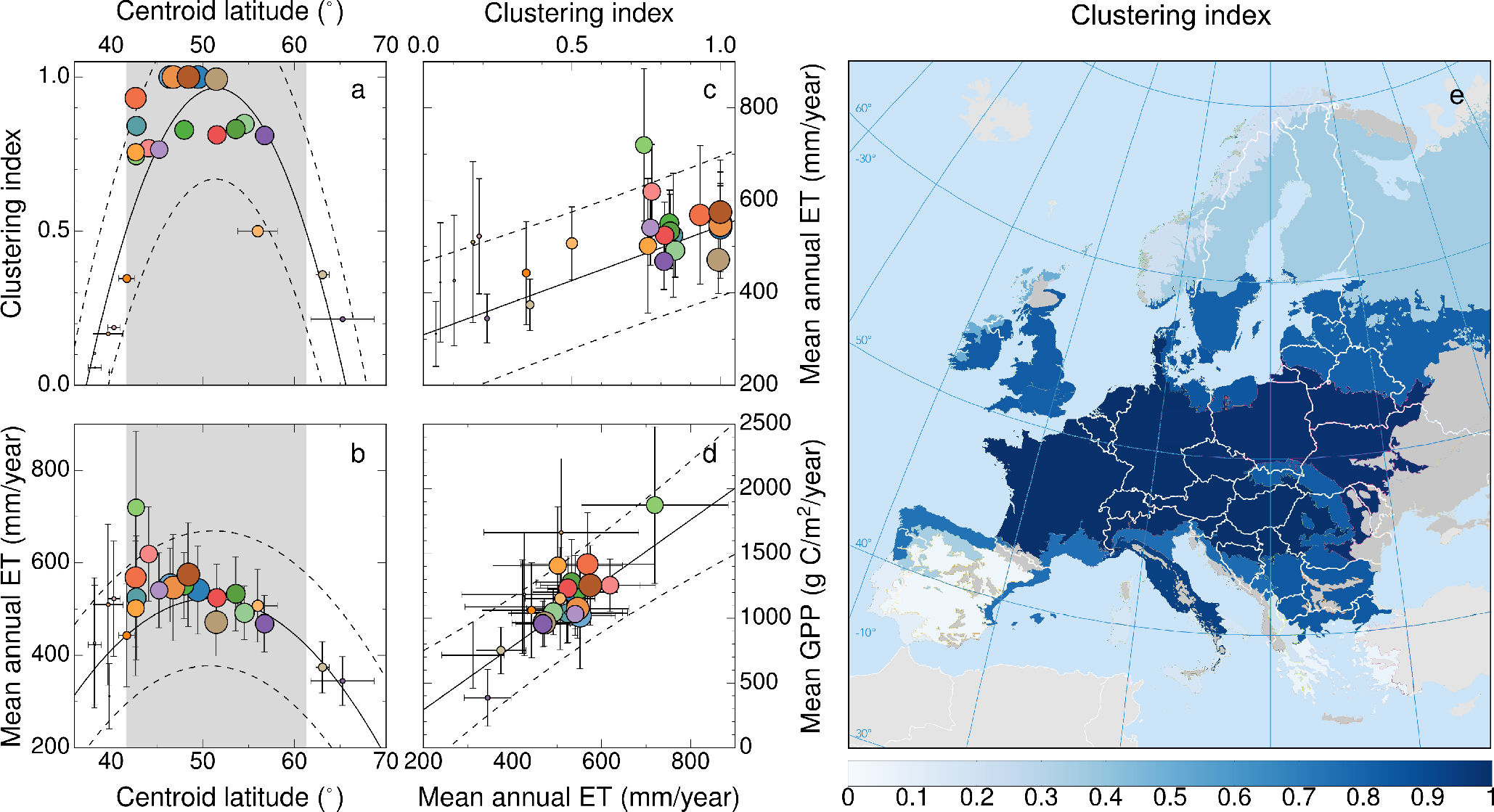
Linking height clustering to geographical and environmental variables. **a**, Variation in the clustering index (*q*) with latitude (φ). Quadratic fit: *r*^2^ = 0.77, *p* < 10^−3^. **b**, Latitudinal variation in mean annual actual evapotranspiration (ET) data. Quadratic weighted regression: *r*^2^ = 0.63, *p* < 10^−3^. The shaded areas in panels **a** and **b** represent the latitudinal range for which the adjusted dependence *q*(φ) ≥ 1/2, where both height clustering and evapotranspiration are maximal. **c**, Linear weighted regression for ET as a function of the clustering index; *r*^2^ = 0.60, *p* < 10^−3^. **d**, Correlation between mean gross primary productivity (GPP) and mean annual ET; linear weighted fit: *r*^2^ = 0.73, *p* < 10^−3^ In the first four panels, the radius of each circle is proportional to the clustering index. Symbol colors refer to different ecoregions (Fig. 1). All the fits show the 95% prediction intervals. **e**, Geographical distribution of clustering indices for ecoregions across Europe.

## Discussion

To the best of our knowledge, this is the first time that predictions from modern coexistence theory [5] and the competition-similarity paradigm [6] have been tested with macro-ecological trait data at large spatial scales. While potential evapotranspiration decreases with latitude, actual evapotranspiration peaks at intermediate latitudes, and is strongly associated with higher levels of local trait clustering. Critically, actual evapotranspiration is positively correlated with gross primary productivity (GPP) across terrestrial ecosystems (see Fig. 4d and [15]), which also peaks at intermediate latitudes across Europe (Supporting Information, Fig. S3). The confrontation of model predictions against plant community data across Europe reveals a clear signature of competition in local communities in the environmentally conducive middle-range latitudes; as environmental conditions get increasingly extreme, they override competitive effects resulting in non-significant clustering in plant heights.

Harsh environments either interfere with competitive interactions or create conditions for other types of interactions to drive community assembly. In either case, plant trait clustering is not apparent. Although competition may play a role, particularly at small spatial scales, facilitation is likely to be more important under stressful conditions [38]. By contrast, the mild environments characteristic of mid-latitude ecoregions impose less stringent limits to plant growth and the effect of species interactions through competitive dominance can be observed even at large biogeographic scales. At these intermediate latitudes, differences in competitive ability tend to be weaker, on average, and trait clustering is likely to arise from competitive interactions. In these ecoregions, competition, by filtering out subdominant species, leaves a significant trace on local community assembly and, as empirical data confirm, species tend to cluster around smaller height values (Supporting Information, Fig. S8). According to the conceptual framework that explains plant height as a trade-off between light and water availability, the over-representation of small heights is interpreted as a fingerprint that accessing water resources could influence plant growth more importantly than accessing light at mid-range latitudes. Interestingly, the relevant role of competitive dominance driven by species trait hierarchies has been also reported at much smaller spatial scales for forest trees in the French Alps [12]. In addition, a recent study of the assembly of forest communities across East Asia shows that a phylogenetic-based species similarity index tends to be smaller the higher the minimum temperature of the coldest month is [39]. Together with our results, these studies suggest that trait clustering is generally likely to occur where conditions for plant growth are less restrictive. Our model indicates that the process underlying this pattern is competitive dominance, although it is likely that community assembly for other taxa may be driven by other biotic or environmental filters. For instance, phytoplankton communities appear to be driven by limiting similarity creating clumpy species coexistence in estuarine ecosystems [40].

This analysis uncovers a new macro-ecological empirical pattern involving the relationship between trait clustering in maximum stem height of plant species and plant primary productivity (and actual evapotranspiration) over large spatial scales. The intensity of this clustering increases with primary productivity, this is, it is more significant in regions with better conditions for plant growth. Conversely, in harsh environments this clustering disappears. Our theoretical investigations point to the role of competitive dominance as a plausible explanation underlying this pattern at the local level. The relevance of this mechanism at driving community assembly, which tends to equalize differences in competitive ability, should increase with gross primary productivity. Our strategy provides a clear direction for quantitatively assessing the generality of this prediction in other regions, for different taxa, and for other traits.

Theoretical approaches to understand the forces shaping ecological communities rely on mathematical models of species interactions in order to predict community assembly rules, species coexistence, and community stability. Empirical approaches rely, instead, on sampling existing ecological communities to collect as much information as possible on species composition, abundances, and associated environmental variables across temporal and spatial scales in order to infer regularities using statistical models. Finding a theoretically robust and ecologically meaningful rapprochement between these longtime independent approaches at relevant scales remains a challenge for ecology, and we trust that our work will inspire new contributions in this direction.

## Methods

Additional information on methods is provided as a single Supporting Information document containing notes clarifying statistical analyses, a full assessment of the robustness of the results, and a detailed description of the stochastic community model.

## Plant species data

Mean height values were obtained from the LEDA database [28] for as many species as there were available in that source. Missing values were taken from [41]. According to plant growth forms, 2610 species were grouped as herbaceous (aquatic, herbs, or graminoid) and 623 as woody (shrub or tree). Here we reported results for herbaceous plants, although similar conclusions were obtained for the entire data set (see Figs. S5, S6, and S7 in Supporting Information). Height values spanned several orders of magnitude, so we used a log-transformed variable (*h* = log *H*) to measure species differences (using non-transformed heights yielded comparable results, see Supporting Information, Figs. S5, S6, and S7). The values of h were standardized within ecoregions as *t* = (*h* — *h*_min_)/(*h*_max_ — *h*_min_) so that 0 ≤ t ≤ 1.

In a metacommunity with richness *S*, a number *s_k_* ≤ *S* of species will form a local community assemblage at cell *k*. The coexistence probability is defined as the average fraction 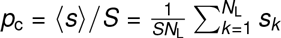, with *N*_L_ representing the number of cells in the ecoregion. This quantity, together with the distribution of trait differences in cells, was used to compare model predictions with real data.

## Stochastic community model

Community dynamics is mathematically described as a birth-death-immigration process in continuous time (Supporting Information, Sec. S2.2). Let **n** = (*n*_1_,…, *n_S_*) be a vector of population numbers for all species in the metacommunity. At each time step, one of the following events can take place: (1) immigrants arrive from the metacommunity at rate *µ*, (2) local births and deaths occur at rates *α*^+^*n_i_*, and *α*^−^*n_i_*, respectively, for species *i*, (3) two individuals of the same species *i* compete at a rate 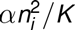, where *α* = |*α*^+^ — *α*^−^|, (4) two individuals of distinct species *i* and *j* compete at a rate *αρijn_i_n_j_*/*K*, resulting in an increase of population size n_i_ if *ρij* < 0 and a decrease if *ρij* > 0 (Supporting Information, Sec. S2.1). For the sake of simplicity, in simulations trait values were drawn from a Gaussian distribution with zero mean and variance *σ*^2^. Trait differences were transformed to interaction strengths as 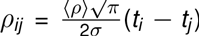. With this choice, the expected value of |*ρij*| is precisely 〈*ρ*〉, and we simply varied 〈*ρ*〉 in model simulations to produce the curves shown in Fig. 3a (Supporting Information, Sec. S2.1).

## Randomization tests

Our randomization tests were based on the average competitive strength observed in a local cell formed by *s* species, (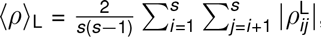, where 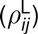 is the submatrix of the metacommunity competition matrix restricted to the species present in the local community. Compared to metacommunity samples, the lower (higher) the empirical local average 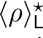 is, the higher (lower) is the degree of species clustering in the local community. For each local community we calculated the probability 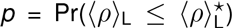 that the empirical local average is smaller than the randomly-sampled competition average. At a 5% significance level, if *p* > 0.95 the empirical competition average is significantly larger than the average measured for random metacommunity samples, which implies that average trait differences in local communities are larger than expected at random. On the other hand, if *p* < 0.05, observed trait differences are significantly smaller than expected at random. Therefore, if *p* > 0.95, the local community exhibits ‘significant trait overdispersion’, whereas if *p* < 0.05, there is evidence for ‘significant trait clustering’ in the local assemblage.

## Acknowledgements

The authors thank Mercedes Pascual and Stefano Allesina for their comments, and are indebted to Rohan Arthur for his constructive criticism of the manuscript. This work was funded by the Spanish ‘Ministerio de Economia y Competitividad’ under the project CGL2012-39964 (DA, JAC) and the Ramon y Cajal Fellowship program (DA). JAC acknowledges partial financial support from the Department of Applied Mathematics, Technical University of Madrid.

## Author contributions

JAC and DA conceived and designed the research; JAC and SC performed the statistical analysis and model simulations; JAC, SC, AO and DA analyzed the data and results; AO contributed materials; JAC and DA wrote the paper.

